# Significantly improving the quality of genome assemblies through curation

**DOI:** 10.1101/2020.08.12.247734

**Authors:** Kerstin Howe, William Chow, Joanna Collins, Sarah Pelan, Damon-Lee Pointon, Ying Sims, James Torrance, Alan Tracey, Jonathan Wood

## Abstract

**Background:** Genome sequence assemblies provide the basis for our understanding of biology. Generating error-free assemblies is therefore the ultimate, but sadly still unachieved goal of a multitude of research projects. Despite the ever-advancing improvements in data generation, assembly algorithms and pipelines, no automated approach has so far reliably generated near error-free genome assemblies for eukaryotes.

**Results:** Whilst working towards improved data sets and fully automated pipelines, assembly evaluation and curation is actively employed to bridge this shortcoming and significantly reduce the number of assembly errors. In addition to this increase in product value, the insights gained from assembly curation are fed back into the automated assembly strategy and contribute to notable improvements in genome assembly quality.

**Conclusions:** We describe our tried and tested approach for assembly curation using gEVAL, the genome evaluation browser. We outline the procedures applied to genome curation using gEVAL and also our recommendations for assembly curation in an gEVAL-independent context to facilitate the uptake of genome curation in the wider community.

## Background

### Assembly curation adds significant value

Despite the advances in sequencing and mapping technologies and the ever-increasing number of sophisticated algorithms and pipelines available, generating error-free eukaryotic genome assemblies in a purely automated fashion is currently not possible [1,2]. Assembly software designed to generate continuous sequence from raw reads is confused by heterozygous or repeat-rich regions, introducing erroneous duplications, collapses and misjoins. The same issues recur in subsequent scaffolding processes that aim to turn primary contigs into representations of chromosomal units. The fact that these tools are commonly applied in series rather than in parallel results in the passing of mistakes made from one process on to the next. As a result, even so-called high-quality or “platinum” assemblies can suffer from hundreds to thousands of duplications, collapses, misjoins and missed joins. Because assemblies are often judged simply by their continuity, rather than by their completeness and (structural) correctness, these errors go unnoticed. This impacts research in many ways, making whole regions of the genome impossible to access or misleading researchers who misinterpret assembly artifacts as biological findings [3]. One way to address these shortcomings is in-depth analysis of discordances between the assembly that has been generated and the different data types available for the sequenced individual or species and subsequent resolution of these discordances. This can be performed at the sequence and the structural level. Many automated tools are available that assess sequence quality through read alignment, kmer counting, gene finding and other methods [4–7]. For structural quality assessment, several individual tools can be used, but these tend to analyse a single data type at a time rather than combining insights from analysis of several in parallel [8,9].

We created gEVAL, the genome evaluation browser, to enable a user to visualise and evaluate discordances between an assembly and multiple sets of accompanying data at the same time [10]. gEVAL enables the identification of errors and simultaneously suggests ways to resolve them. Combined with manual assessment of the generated data by experienced curators and a pipeline that enables the curators to record changes and recreate the improved assembly accordingly, gEVAL provides a critical addition to strategies striving to produce assemblies of the highest possible quality.

Below we outline the strategic design, achievements and limitations of the gEVAL approach to assembly curation. gEVAL is tied into our local infrastructure and as such sadly not portable. We therefore also provide detailed recommendations on how to create similar analyses that do not use gEVAL to promote the core, proven design concepts in gEVAL.. This is especially timely in the context of emerging projects that aim to assemble the genomes of very large numbers of species to highest quality possible, including the Vertebrate Genomes Project (VGP), the Darwin Tree of Life Project (DToL, darwintreeoflife.org) and the overarching Earth Biogenome Project (EBP) [1,11].

## Results

### Checking for assembly coherence, coverage and contamination

We recommend that every genome assembly is checked for coherence. This includes making sure that only data that belong to the relevant species are used for assembly in the first place. This is best checked before starting the assembly process by aligning all raw datasets with e.g. mash [12] and checking that the data are in fact combinable (i.e. that they are likely to derive from the same underlying distribution of sequence). A major source of remaining technical error in assemblies is the retention of duplicated regions that result from failure to recognise that two sequences are in fact allelic. These false duplications have wide-ranging negative consequences for subsequent research, for example causing prediction of erroneous gene duplications [1]. False duplications are caused by either incorrect resolution of assembly graphs or failures in detection of haplotypic variation. They can be detected using simple read coverage plots or more sophisticated kmer analyses (for example using KAT, the K-mer Analysis Toolkit [5], KMC, the K-mer counter [13] or Merqury [7]). Kmer approaches also support the estimation of the completeness of the assembly (i.e. whether the assembly contains all the relevant kmers present in the reads) and the ploidy of the genome [14]. False duplications can be removed, ideally after generating the contigs, with tools that recognise partial and complete allele overlap, such as purge_dups [15]. In addition to duplications, assembly quality is also negatively affected by erroneous sequence collapses, mostly located in repetitive regions. Collapses are relatively easy to detect based on increased read coverage, but harder to resolve as they require generation of new sequence. This can be performed through extraction of mapped reads and local reassembly under more stringent conditions, or with more sophisticated methods like SDA [16]. Assemblies are frequently polished after contig generation, using available read data or particular high base accuracy data such as Illumina short reads, to correct remaining errors in the derived consensus sequence. It is however possible to over-polish, such that rare repeat variants are replaced by the most abundant version, or where nuclear insertions of organellar genome fragments (nuclear mitochondrial transfers, NUMTs, and nuclear plastid transfers, NUPTs) are polished to match the organelle sequence. For polishing the target genome assembly therefore must include the organelle genomes. Organellar genomes are often missing from assemblies because assembly toolkits recognise and exclude them as repeat sequence, or because they yield complex graphs that conflict with nuclear insertions. They can be assembled independently, e.g. using the mitoVGP pipeline [17].

Contigs/scaffolds that represent the organelle genomes should be identified and processed independently of the primary, nuclear assembly.

A preliminary assembly of data from a target species can inadvertently include synthetic sequence from cloning or sequencing systems, contamination from species handled in the same laboratory or sequencing centre, or contamination from natural cobionts of the target (the gut and skin microbiomes, unsuspected parasites, etc.). Decontamination serves to detect and mask or remove sequence not originating from the target species, and to separate organelle genomes from the primary assembly if not carried out previously. This includes identifying remaining vector and adapter contamination based on known sequence. Contaminating sequence can be detected with dedicated toolkits, such as BlobToolKit [18] or Anvi’o [19] or through individual sequence similarity searches using BLAST or Diamond against suitable databases (Table 1). Our in-house pipelines use automated detection of synthetic, laboratory and natural contaminants, but include manual controls to preserve sequences that may be the product of horizontal gene transfer (described below).

**Table 1:** Detecting decontamination in assemblies, inspired by the processes carried out by GenBank’s genome archive [20].

| *Detection of* | *Software tools* | *Detection requirements* | *Database* |
| --- | --- | --- | --- |
| vector/adaptor<br>sequence | Vecscreen [21] |  | UniVec [22] |
| common<br>contaminants | megaBLAST [23] | e-value $\leq 1e-4$, reporting matches $\geq 98\%$<br>sequence identity with match length 50-99 bp,<br>$\geq 94\%$ with match length 100-199 bp, or $\geq 90\%$ with match length $> 200$ bp | Contamination in<br>eukaryotes [24] |
| organelle genomes | megaBLAST | e-value $\leq 1e-4$, sequence identity $\geq 90\%$,<br>match length $\geq 500$ | RefSeq mitochondria [25]<br>and plastid [26]<br>assemblies |
| other species | megaBLAST | e-value $\leq 1e-4$, match score $\geq 100$, sequence<br>identity $\geq 98\%$; ignore regions also matching<br>highly conserved rDNAs | Windowmasked [27]<br>RefSeq genomes [28] |

Lastly, trailing Ns should be removed from all contigs and scaffolds.

### Improving structural integrity

As most assembly pipelines currently apply different scaffolding steps in series, errors in early steps can propagate through the process. To avoid compounding these errors, one could carry out a thorough curation process after every scaffolding step, but if many scaffolding steps are involved this will be very demanding on time and resources. Our experience has shown structural integrity can be successfully improved after completion of a full, automated assembly process [1,10].

The principle behind identification of assembly errors is simple: align all available (raw and other) data to the produced assembly, check for discordances, and then correct. Several tools that detect scaffolding issues with single data types are available, including scaff10x for 10X Chromium linked reads [29], Access for BioNano maps [8], and HiGlass [30], pretext [31] and Juicebox [9] for Hi-C data. ASSET evaluates multiple data types in parallel [32]. Read coverage plots identify errors or problem regions through deviation from expected averages (indicating possibly problematic low-coverage regions, haploid regions, or regions of collapsed repeat) and sites where aligned reads are all clipped at the same site (suggesting that the assembly contains an erroneous join). Aligning the assembly against itself can be used to detect duplications.

Additional data not used in generating an assembly also provides critical information. Comparing the assembly to previous assemblies from the same species or to assemblies from closely-related species can highlight areas of disagreement, and thus areas that deserve closer attention during curation. Transcript evidence, such as assembled cDNAs or long single-molecule reads, can be aligned to affirm joins across sequence gaps, identify local mis-assemblies, and to detect false duplications. Protein sequences from the same or related species can serve the same purpose. Centromeres and telomeres can be identified in the assembly through sequence features [33,34]. Long-range structural data (such as karyotypes and FISH mapping) and genetic mapping data (such as meiotic mapping or radiation hybrid mapping data) can provide validation of the large-scale correctness of an assembly, and in particular guide correct association and orientation of chromosomal arms with respect to telomeres and centromeres. Chromosome-wide patterns of repeat proportion and GC content can also be used to affirm completeness of chromosomal units. Once identified, errors should be corrected. We have found that whole genome sequence editing tools such as gap5 [35], are very useful for this process. It is critical to record the corrections made so that the path from primary assembly to the final completed genome assembly is clear and justified.

### Identifying and naming chromosome-scale scaffolds

The ultimate goal of genome assembly is the production of fully contiguous nucleotide sequences that represent each of the chromosomal units for the species, with an estimate of both overall and local quality, and with known sites that may have issues flagged. Long-range data, such as Hi-C contact maps, can reliably indicate which scaffolds correspond to chromosomal units, and these putative chromosomal assemblies can be reconciled with karyotypic information where available. Fully resolved chromosomal units (where all contigs and scaffolds are ordered and oriented) can be submitted to the the INSDC sequence archives (the International Nucleotide Sequence Database Collaboration (INSDC) partners: GenBank, ENA and DDBJ) as a “chromosome”. Scaffolds and contigs that are demonstrably associated with a chromosomal unit but which cannot be joined because of ambiguous order or orientation must be submitted as “unlocalised” for this chromosome. Scaffolds and contigs that cannot be associated with a chromosome, and which also cannot be established as being separate chromosomes, are deemed “unplaced”.

If a reference assembly for the same species or a karyotype with sequence-based anchors is available, chromosome naming should follow the precedent to ensure compatibility with previously reported results. Identification of sex chromosomes can be based on comparisons to related species or from the location of marker genes. In heterogametic individuals, sex chromosomes will also be easily recognisable by their halved sequence coverage compared to autosomes. If no reference for chromosome naming is established, they should be named by size.

Last but not least, every assembly, together with all relevant raw and metadata, should be submitted to one of the INSDC archives (Genbank, ENA or DDBJ, [36]) to allow discoverability, assure community access and provide stability for future analyses.

### Assembly curation for high-throughput projects

The above described curation processes suffer from the same shortcoming as the assembly process itself: they are usually applied in series rather than in parallel. The benefits of a multitude of data types and approaches are also difficult to realise. Whilst the identification of many assembly issues can be automated, the actual decision to apply a change is still best made by an experienced curator, seemingly slowing the process to an extent that excludes it from any high-throughput project.

The Genome Reference Informatics Team (GRIT) assembly curation pipeline was established to deliver high quality assembly curation for the Genome Reference Consortium (GRC, [37]), the VGP and DToL. The pipeline automates the processes of gata gathering and computational analysis for decontamination, validation and correction of assemblies, sourcing all available data from in-house and public resources. The analyses are then presented for manual evaluation by experienced genome curators, who perform the evaluation and log required changes. The corrected assembly ready for submission is generated automatically. Central to this pipeline is gEVAL, the genome evaluation browser [10]. gEVAL enables visualisation and evaluation of discordances between an assembly and multiple sets of accompanying data in parallel, enabling the simultaneous identification of errors and ways to resolve them [38]. The pipeline that GRIT deploys is closely tied into the Wellcome Sanger Institute’s internal data structure and cannot be ported, but is described here as an example of a successful implementation that mixes automated and manual processes and significantly improves genome assemblies in a time and resource sensitive way that allows its use within high-throughput projects.

The GRIT curation process usually starts with assemblies that have been purged of duplicates and most haplotypic segments, scaffolded with long-range data and polished. Before being loaded into gEVAL, all assemblies are run through a nextflow [39] pipeline that performs contamination detection and separation or removal as described in Table 1, combined with removal of trailing Ns [39]. Brief manual checking of the results prevents the erroneous removal of regions likely derived from horizontal gene transfer.

gEVAL analyses are collated in a database built on an Ensembl framework [40] that has been modified to visualise assembly quality rather than gene and feature annotation. Loading of the analyses into gEVAL and subsequent assembly analyses are pipelined using snakemake and vr-runner [41,42]. Which analyses are run and visualised depends on the availability of data, but typically include the types listed in Table 2. The alignments and placements are visualised in a genome browser as feature tracks and colour-coded to indicate agreement or disagreement with the assembly (Fig. 1). The gEVAL process also generates lists that detail discordances between the assembly and the different data types. The process of analysis and loading into gEVAL requires up to 3 days for a 1 Gb assembly.

**Table 2:** Examples of data types and analyses included in gEVAL and their ability to detect issues and errors.

| *Data type* | *Software* | *Supports analysis of* |  |  |  |
| --- | --- | --- | --- | --- | --- |
|  |  | *misjoins* | *missed joins* | *duplications* | *collapses* |
| long reads | Minimap2 [43] ,<br>winnowmap [44] | x | x | x | x |
| BioNano | BioNano Solve | x | x | x | x |
| 10X linked reads | Break10x [29] | x |  |  |  |
| cDNAs/gene sets | Blat [45] pblat [46] | x | x | x |  |
| self-alignments | Mummer [47] |  |  | x |  |
| other assemblies | Compara [40] | x | x |  |  |
| centromeres | Repeatmasker [40,48]<br>centromere db [33] | x | x |  |  |
| telomeres | Find_telomere [34]<br>adapted to work with any<br>sequence | x | x |  |  |
| genetic and other maps | EPCR [49], Blast [50] | x | x | x |  |

**Figure 1:**
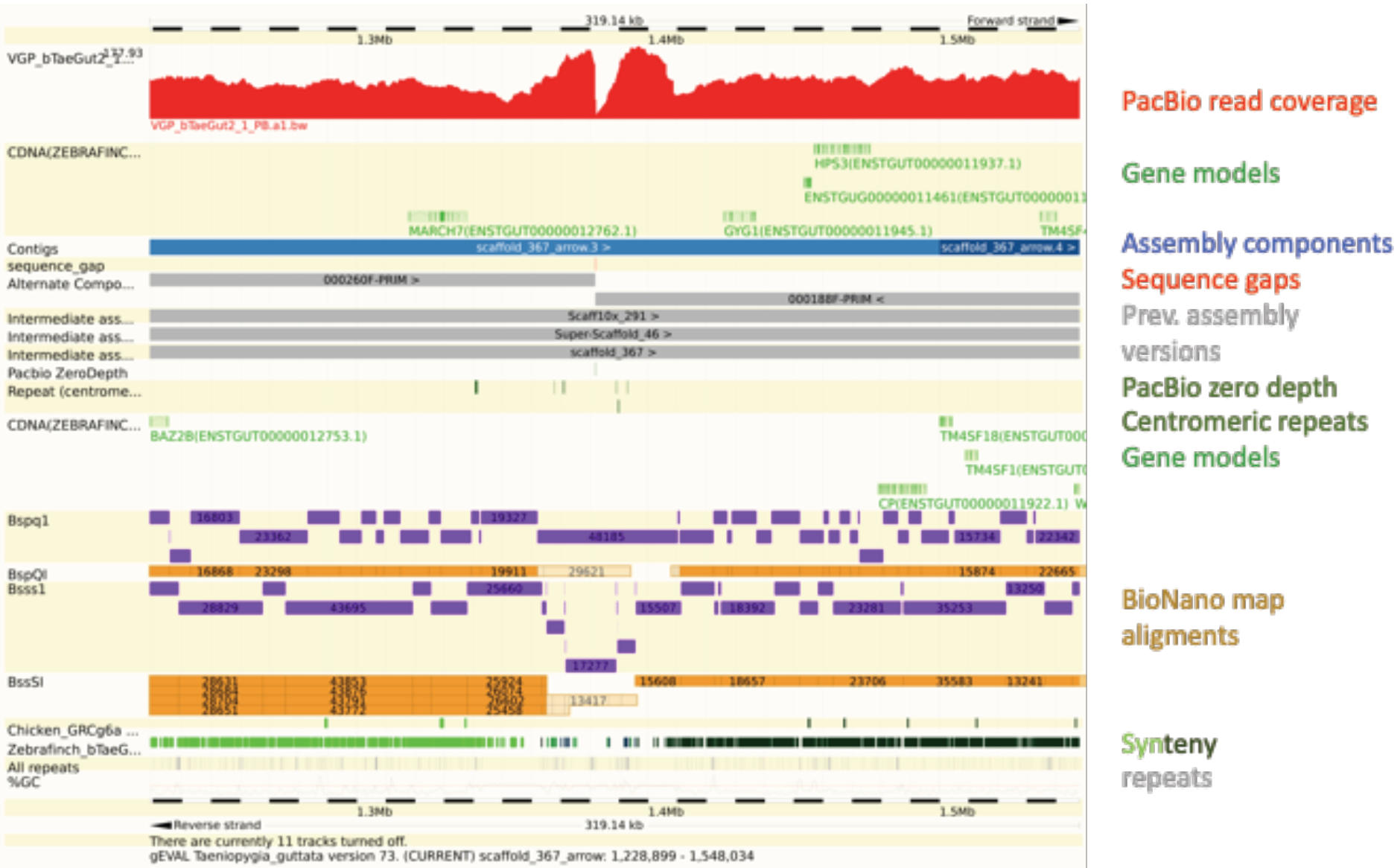
Example of an assembly issue identified in gEVAL in a bird genome (*Taeniopygia guttata*, VGP). Feature tracks (named on the right) are shown in the context of the assembly. A misjoin is visible in the middle of the example, indicated by the drop in Pacific Biosciences read coverage, discordance with the aligned BioNano maps and the break in synteny. The alignments with intermediate assembly stages show that this error was introduced by the scaffolding step involving scaff10x.

gEVAL automatically flags areas where the raw and other comparative data available are discordant with the presented assembly. Experienced curators use the gEVAL database and visualisation, and (where available) Hi-C maps (generated outside the gEVAL pipeline and viewed in HiGlass [30], or pretext [31]), to check each listed discordance and decide whether and how to adjust the sequence based on the available data. In rare cases, the information contained in gEVAL and the Hi-C maps is not sufficient to decide whether a change is warranted. The curators then use additional tools such as gap5 [35] for in-depth analysis of aligned reads or Genomicus for information on synteny with other species [51]. Curators propose a variety of interventions such as breaking or joining sequence regions, changing the order and orientation of scaffolds and contigs and removing false duplications. Detangling sequence collapses is currently only possible where additional data can be employed for local reassembly. In high-throughput projects such as DToL or VGP curation is usually restricted to a resolution of around 100 kb. This allows an experienced curator to complete curation of 1 Gb of sequence in around 3 days. For projects without immediate time restraints and aimed at single references, such as the genomes curated within the GRC, there is no resolution limit.

During the gEVAL build, assembly scaffolds are split into equally sized components, with their order and orientation recorded in a path file under version control, listing component name, scaffold name and orientation. Should any rearrangement be necessary, the curators simply reorder/reorient the components in the path file. If necessary, components can be split with bespoke scripts which create new components and store them in the gEVAL database. After manual curation, the adjusted ordering and orientation of components and a list of scaffold-chromosome associations is processed automatically to generate the final assembly for submission. All milestones and metrics of the whole curation process are recorded in a tracking database.

### Using gEVAL to assess published assemblies

Above we have described the use of gEVAL to create high-quality assemblies. gEVAL can also also be used to support research communities in verifying research results, ensuring they are not based on assembly artifacts. For this, a gEVAL database is generated for publicly available assemblies, as e.g. is the case for all GRC assemblies [38]. Here, gEVAL offers the same analyses as detailed above, plus additional databases with other assemblies of the same species, such as previous versions of the current reference, including whole genome alignments between them (Fig. 2). Combined with tutorials and documentation, this provides a valuable resource for users of the featured reference assemblies.

**Figure 2:**
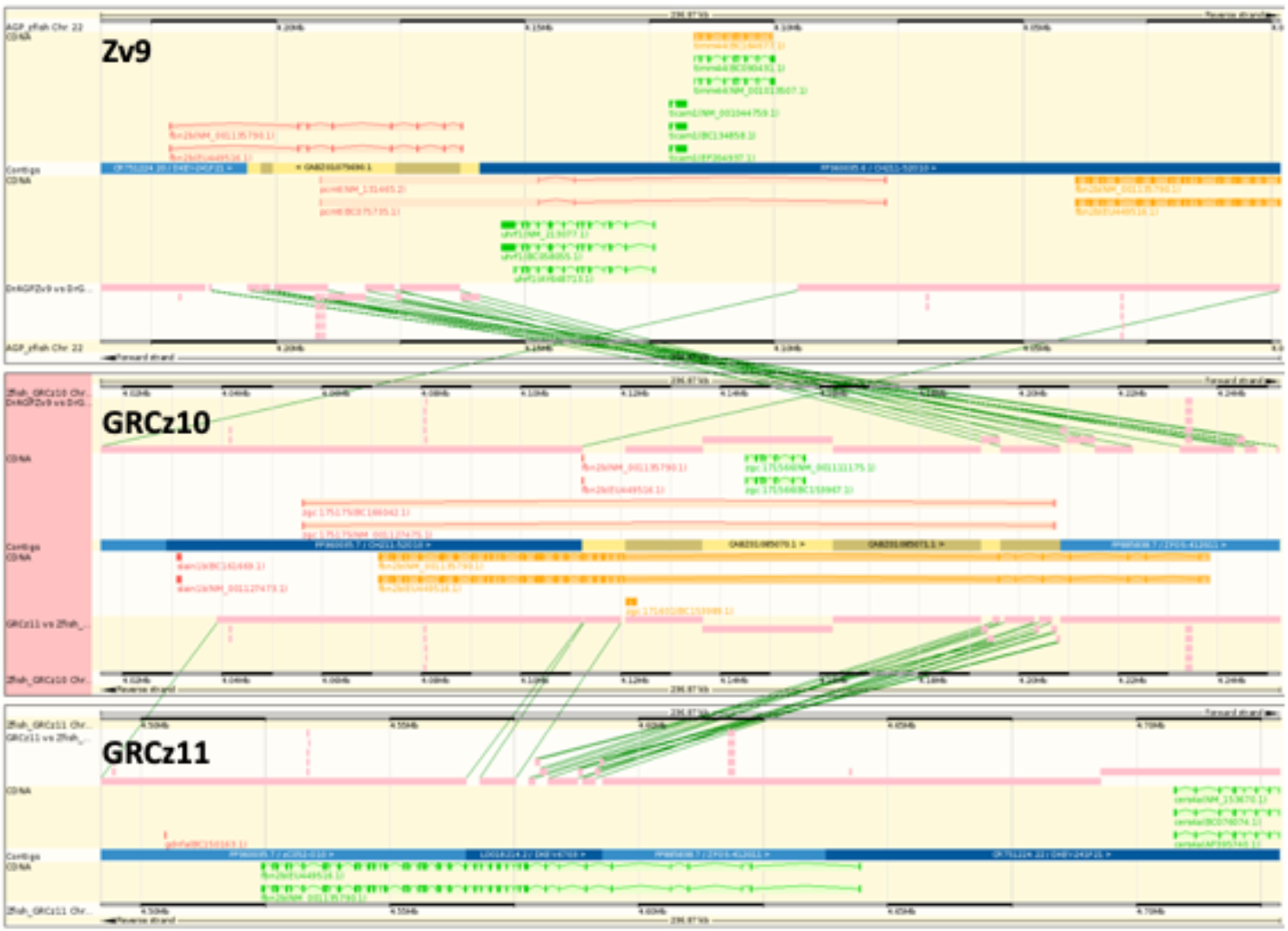
Comparison of the *fbn2b* region in the *Danio rerio* (zebrafish) reference assemblies Zv9 (top), GRCz10 (middle) and GRCz11 (bottom) in gEVAL. The fragmented *fbn2b* locus (colour coded in orange and red) was adjusted for GRCz10 (colour coded in orange) and further improved by removing whole genome shotgun contigs in favour of finished clone sequence for GRCz11. The final correct gene locus is coloured in green.

### Impact of assembly curation for high throughput-projects

During curation of 111 assemblies (174 Gb sequence) for VGP and DToL, on average 221 interventions per Gb of sequence were applied (67 breaks, 105 joins and 49 removals of false duplications, Fig. 3). These changes led to an average reduction in assembly length by 2% as the curation effort did not generate new sequence. However, average scaffold N50 increased by 40% and scaffold number decreased by 29%. It is important to note that scaffold N50 changes differed for each assembly, and that while the process improved N50 several hundred-fold in initially fragmented assemblies it (up to) halved the N50 in over-scaffolded assemblies. On average 96% of assembly sequence was scaffolded to chromosome-level (Fig. 4).

**Figure 3:**
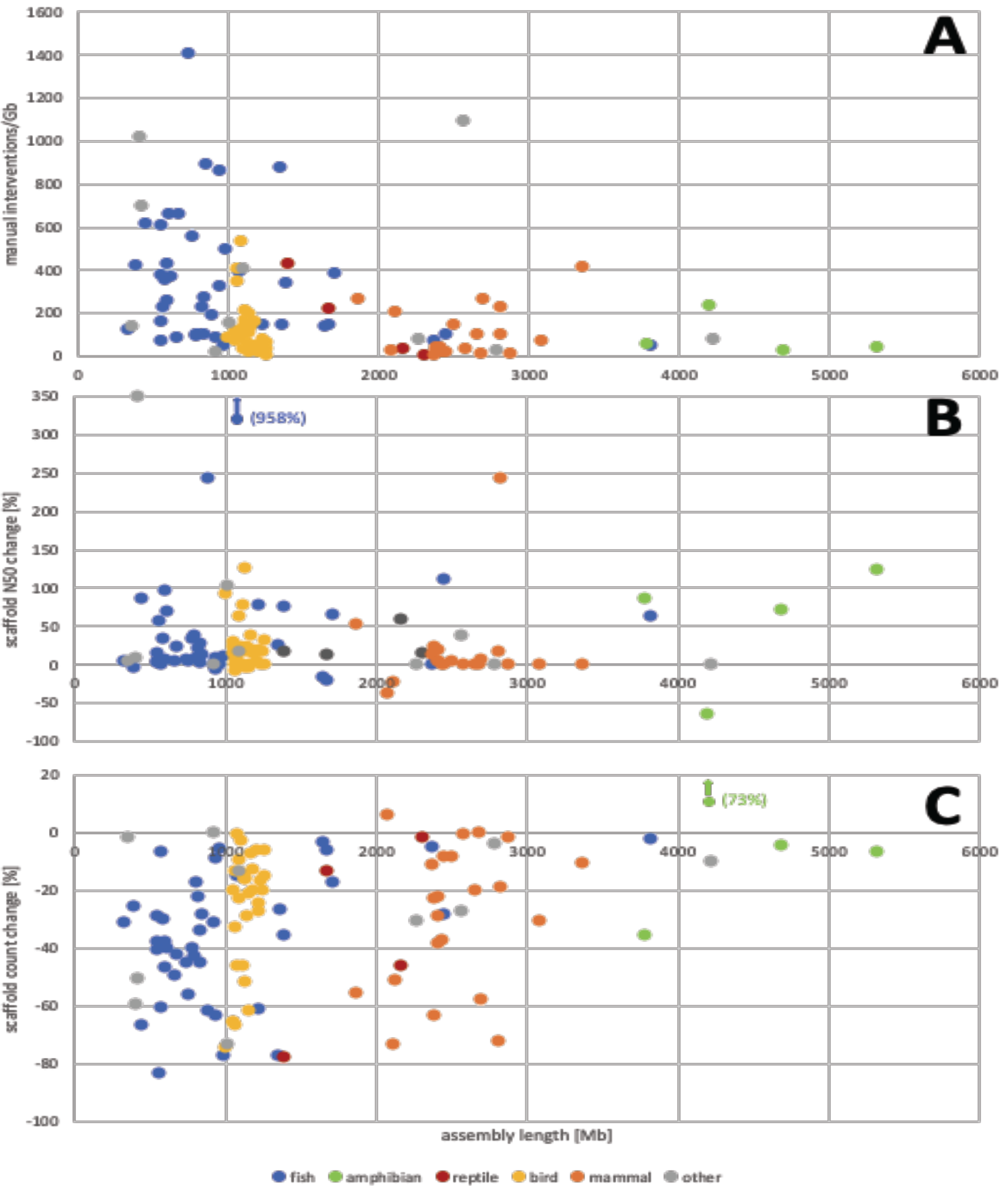
Changes to 111 assemblies from different clades through manual assembly curation by the Genome Reference Informatics Team at the Wellcome Sanger Institute. (**A**) Manual interventions (breaks, joins, removal of false duplications) as events per Gb of assembly sequence. (**B**) Changes in scaffold N50 after curation. (**C**) Changes in scaffold counts after curation. The depicted assemblies were created with PacBio CLR, Chromium 10X and Hi-C data, most also included BioNano data.

**Figure 4:**
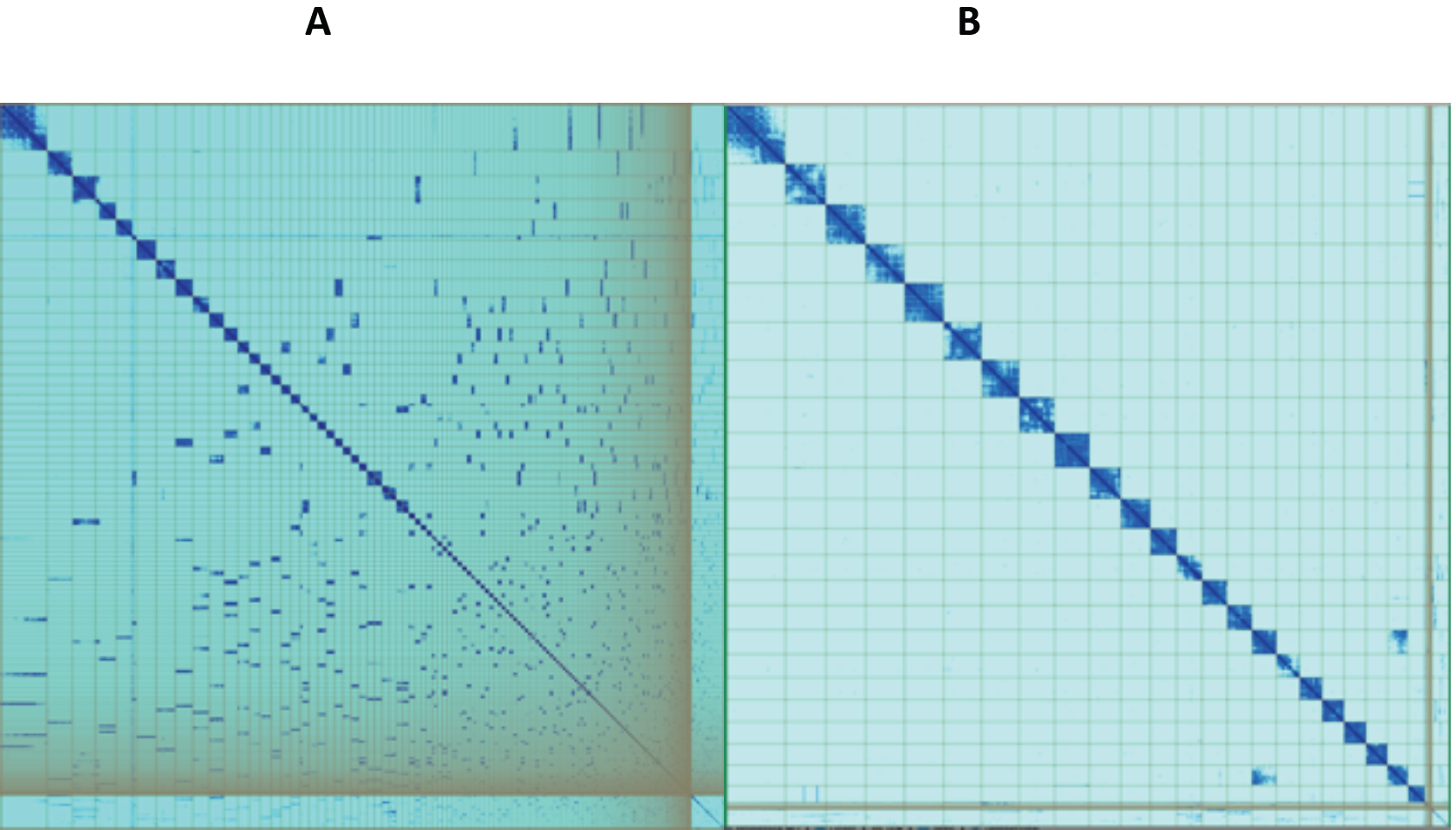
Hi-C maps (pretext) showing the *Asterias rubens* (starfish) genome assembly (sequenced as part of the Sanger Institute’s 25 genomes for 25 years project) before (A) and after (B) curation. The curation corrected the initial assembly by making 75 breaks and 216 joins and removed one stretch of erroneously duplicated sequence. 97% of the assembly sequence could be assigned to 22 chromosomes. The curated assembly (**B**) contains one scaffold that is known to be associated with a second one (off-diagonal signal at bottom right), but its order and orientation are ambiguous. This scaffold has been submitted as “unlocalised” for the relevant chromosome.

## Conclusions

The number and scale of changes to the assemblies necessary across the diversity of species analysed shows the persistent need for manual intervention on the path to high quality genome assemblies. Our experiences in curating genomes for GRC, VGP and DToL have driven improvements in assembly software (e.g. purge_dups [15], salsa2 [52]), assembly pipelines (VGP, DToL) and assembly assessment tools (e.g. Asset [32]). Genome assembly generation is a fast-moving field and we are constantly adapting the curation software and processes to include novel data types whilst being conscious of the need to maximise throughput. This ensures ongoing involvement of assembly curation in high-throughput projects to produce the best possible data for the community to base their research upon.

## Authors’ contributions

KH conceived the project and drafted the manuscript. WC implemented gEVAL, WC and YS implemented the curation pipeline. JT performed contamination checking, compartimentalisation and assembly postprocessing. WC, YS, JW and DPL built gEVAL databases; JC, SP, AT and JW curated genome assemblies.

## Acknowledgements

We thank Mark Blaxter, Richard Durbin and Shane McCarthy for their input on this project and many helpful discussions. Additional thanks go to Mark Blaxter for many constructive discussions around this manuscript. The genome curation project is heavily influenced by the Genome Reference Consortium, the Vertebrate Genomes Project and the Darwin Tree of Life Project and we are indebted to all the members for their engagement with the curation process.

